# Regulatory coiled-coil domains promote head-to-head assemblies of AAA+ chaperones essential for tunable activity control

**DOI:** 10.1101/166256

**Authors:** Marta Carroni, Kamila B. Franke, Michael Maurer, Jasmin Jäger, Ingo Hantke, Felix Gloge, Daniela Linder, Sebastian Gremer, Kürşad Turgay, Bernd Bukau, Axel Mogk

## Abstract

Ring-forming AAA+ chaperones exert ATP-fueled substrate unfolding by threading through a central pore. This activity is potentially harmful requiring mechanisms for tight repression and substrate-specific activation. The AAA+ chaperone ClpC with the peptidase ClpP forms a bacterial protease essential to virulence and stress resistance. The adaptor MecA activates ClpC by targeting substrates and stimulating ClpC ATPase activity. We show how ClpC is repressed in its ground state by determining ClpC cryo-EM structures with and without MecA. ClpC forms large two-helical assemblies that associate via head-to-head contacts between coiled-coil middle domains (MDs). MecA converts this resting state to an active planar ring structure by binding to MD interaction sites. Loss of ClpC repression in MD mutants causes constitutive activation and severe cellular toxicity. These findings unravel an unexpected regulatory concept executed by coiled-coil MDs to tightly control AAA+ chaperone activity.

## Introduction

AAA+ (ATPase associated with a variety of cellular activities) proteins control a multitude of essential cellular activities including DNA replication and recombination, protein transport and quality control, among many others ^1^. AAA+ proteins convert the energy derived from ATP hydrolysis into a mechanical force to remodel bound substrates. ATP binding and hydrolysis is mediated by the conserved AAA domain, which also mediates AAA+ protein oligomerization typically into hexameric assemblies harboring a central channel. Substrate remodeling involves a threading activity into the central channel mediated by mobile loops that bind substrates and pull them upon nucleotide-induced motions ^2,3^. AAA+ proteins gain functional diversity by extra domains that are either fused to or inserted into AAA domains. Extra domains enable targeting of specific substrates but they can also globally control AAA+ protein activity ^4^.

AAA+ proteins are key players in protein quality control by targeting misfolded and aggregated proteins to degrading and refolding pathways. ClpB/Hsp104 reactivates aggregated proteins in concert with a cognate Hsp70 system ^5,6^. Other AAA+ proteins (e.g. ClpX, Rpt1-6) associate with peptidases (e.g. ClpP, 20S proteasome) to form AAA+ proteases, feeding protein substrates into associated proteolytic chambers for degradation ^7,8^. The unfolding activity of AAA+ proteins can, however, also be deleterious to cells, in particular if linked to protein degradation, and therefore needs to be tightly controlled. Accordingly, loss of the control mechanisms of AAA+ protein activity in mutant proteins can lead to cell death ^9-11^.

A high degree of substrate selectivity is equally important to ensure proper functioning of AAA+ proteins and to prevent deleterious activities. Substrate specificity is achieved by adaptor proteins, which bind selective substrates and target them to the AAA+ partner chaperone. Their functions can be additionally controlled by post-translational modifications ^12^, counteracting anti-adaptors ^13,14^ or degradation ^15^. Adaptor proteins can furthermore regulate the ATPase activity of AAA+ proteins and couple substrate delivery to ATPase activation ^9,16^.

Activity control and adaptor action requires the ATPase activity to be repressed in the ground state, which is key to AAA+ chaperone mode of action. Repression can be achieved by regulatory coiled-coil domains inserted into an AAA module. In the ClpB/Hsp104 disaggregase a long coiled-coil middle domain (MD), consisting of two wings, is forming a repressing belt around the AAA ring to reduce ATPase activity ^17,18^. Adjacent MDs bind to each other by head-to-tail interactions keeping the regulatory domains in place. ATPase repression is relieved by MD dissociation and binding to Hsp70 adaptors that prevent reassociation of MDs with the ClpB/Hsp104 ring ^9,19,20^.

The bacterial AAA+ chaperone ClpC associates with the peptidase ClpP to form a central proteolytic machinery of Gram-positive bacteria. The ClpC-ClpP machinery acts in regulatory and general proteolysis, controlling multiple cellular pathways and differentiation processes and is crucial for bacterial stress resistance and virulence ^21-28^. ClpC activity crucially relies on cooperation with adaptor proteins including MecA, that target specific substrates while concurrently stimulating ClpC ATPase activity ^15,16,21,29^. MecA binds to N-terminal and middle domains of each ClpC subunit forming a separate layer on top of the ClpC AAA ring ^30^. How ClpC is kept inactive in adaptor absence, and how the adaptor activates the ATPase is largely unknown. Furthermore, ClpC harbors a coiled-coil MD consisting only of a single wing, as opposed to the two-wing MD of ClpB/Hsp104. A potential regulatory function of the ClpC MD has not been investigated, but its smaller size as compared to the ClpB/Hsp104 MD implies it must act differently if involved in ClpC activity control. Understanding ClpC regulation is particularly relevant as AAA+ protease machines including ClpC have attracted considerable attention as targets for antibacterial action in recent years ^31^. Overruling AAA+ protease control by small molecules can lead to constitutive uncontrolled and toxic activation as best exemplified by acyldepsipeptide antibiotics of the ADEP class targeting the ClpP peptidase. ADEP-activated ClpP causes aberrant protein degradation and even allows for eradication of *Staphylococcus aureus* persister cells ^32-34^. Understanding ClpC activity control therefore might open new avenues for antibiotics development.

Here, we report on an unexpected mode of AAA+ chaperone control involving transition between an inactive resting state and a functional hexamer as revealed by determining the cryoEM-structures of *S. aureus* ClpC in absence and presence of MecA. The ClpC resting state is composed of two helical ClpC assemblies stabilized by head-to-head MD interactions. MecA prevents MD interactions and thereby converts ClpC into a canonical and active hexamer.

## Results

### The ClpC M-domain represses ClpC activity

To study the function of the M-domain (MD) in ClpC activity control we first purified *S. aureus* ClpC/ClpP and demonstrated functionality by determining high-proteolytic activity in presence of the adaptor MecA (**Fig. 1**). Next, we created a series of ClpC MD variants by mutating conserved residues not involved in coiled-coil structure formation (**Supplementary Fig. 1A**). Additionally, we replaced the entire MD (N411-K457) by a di-glycine linker, allowing MD deletion without interfering with folding of the AAA-1 domain. Proteolytic activities of MD mutants were determined using Fluorescein-labeled casein (FITC-casein) as constitutively misfolded model substrate in absence and presence of MecA (**Fig. 1A/B, Supplementary Fig. 1B**). ClpC wild type (WT) together with ClpP exhibited only a low proteolytic activity in absence of MecA and FITC-casein degradation rates were 20-fold increased upon adaptor addition. In contrast, most MD mutants enabled for adaptor-independent FITC-casein proteolysis to varying degrees. ClpC-F436A, ClpC-R443A and ClpC-D444A showed highest activities with degradation rates close to those determined for ClpC WT plus MecA (**Fig. 1A/B**). Similarly, MD deletion strongly increased ClpC activity, indicating that the single point mutants reflect a loss of M-domain function. MecA presence still stimulated FITC-casein degradation by ClpC MD mutants except F436A and ΔM, consistent with the crucial function of F436 in MecA binding (**Supplementary Fig. 1B**) ^30^.

**Figure 1.**
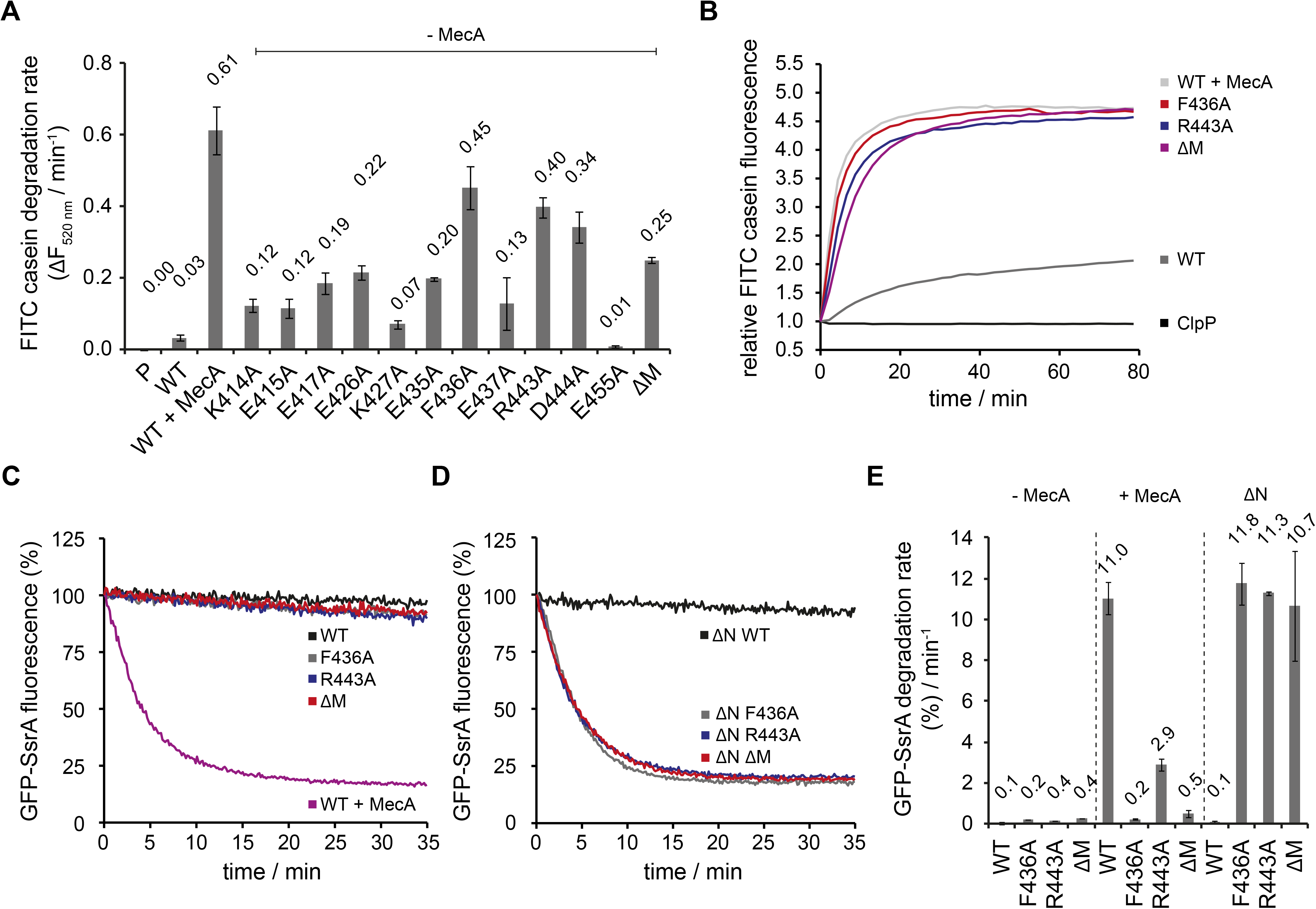
ClpC MD mutants exhibit adaptor-independent proteolytic activity. (**A/B**) FITC- casein degradation was monitored in the presence of ClpP (P) only, or in presence of ClpC wild type (WT, ± MecA) and indicated MD mutants (without MecA). Degradation rates were determined from the initial linear increase of FITC fluorescence. (**C-E**) GFP-SsrA degradation was monitored in the presence of ClpP and indicated ClpC variants. Deletion of the N-terminal domain (ΔN) unleashes high proteolyic activity of MD mutants. GFP-SsrA degradation rates were determined from the initial linear decrease of GFP-SsrA fluorescence.

FITC-casein degradation by activated ClpC M-domain mutants (F436A, ΔM) required ATP hydrolysis and was not observed in presence of ATPγS (**Supplementary Fig. 1C**). Complete degradation of FITC-casein by ClpC-F436A was confirmed by SDS-PAGE (**Supplementary Fig. 2**), while ClpC WT required MecA to exhibit proteolytic activity. Here, we also noticed MecA autodegradation once FITC-casein was digested, in agreement with former findings for the *B. subtilis* ClpC/MecA system ^15,16,29^. We infer that ClpC MD mutants exhibit high, adaptor-independent proteolytic activities, qualifying the M-domain as a negative regulatory element.

To further investigate a repressing function of the ClpC MD we determined GFP-SsrA degradation activities of selected ClpC MD mutants exhibiting highest FITC-casein degradation activities (F436A, R443A, ΔM). GFP-SsrA harbors the 11-meric SsrA tag, which is recognized by AAA+ chaperones pore sites ^35-37^. GFP-SsrA degradation requires application of high unfolding force in contrast to FITC-casein, which is constitutively unfolded. ClpC WT/ClpP efficiently degraded GFP-SsrA in a MecA-dependent manner (**Fig. 1C**). Surprisingly, ClpC MD mutants did hardly exhibit autonomous degradation of GFP-SsrA in contrast to FITC-casein. ClpC-R443A was partially stimulated upon MecA addition demonstrating that the M-domain mutant can process GFP-SsrA in principle (**Supplementary Fig. 3A**). We speculated that differences in GFP-SsrA and FITC-casein binding modes might be the cause of the different degradation activities of ClpC MD mutants. Hsp100 N-domains contribute to casein binding ^38,39^ while their position on top of the AAA-1 ring and central pore site could impede GFP-SsrA binding. We therefore determined GFP-SsrA degradation activities of N-domain deleted ΔN-ClpC and respective MD mutants (**Fig. 1D**). GFP-SsrA remained stable in presence of ΔN-ClpC/ClpP, however, the substrate was rapidly degraded by ΔN-ClpC-F436A, ΔN-ClpC-R443A and ΔN-ClpC-ΔM (+ ClpP) and degradation rates were identical to those determined for ClpC WT/ClpP with MecA (**Fig. 1E**). We did not test for GFP-SsrA degradation by ΔN-ClpC/ClpP in presence of MecA, as the N-domain is essential for MecA binding ^30,40,41^. GFP-SsrA degradation by activated ΔN-ClpC M-domain mutants relied on ATP hydrolysis and remained specific as the substrate variant GFP-SsrA-DD was not degraded (**Supplementary Fig. 3B**). Here, the two C-terminal alanine residues of the SsrA tag are replaced by aspartate residues, obstructing binding to the AAA+ chaperone pore site ^42^. This documents that MD mutations boost ClpC unfolding activity in absence of adaptor without altering general substrate specificity.

### ClpC M-domain mutants exhibit increased basal ATPase activities

*B. subtilis* ClpC activation by adaptors involves strong stimulation of ClpC ATPase activity ^21,29,41^, which we confirm here for *S. aureus* ClpC and MecA (**Fig. 2A**). ClpC WT exhibits a very low basal ATPase activity (0.4 ATP/min/monomer), rationalizing its poor standalone degradation activity. Notably, all activated MD mutants (F436A, R443A, ΔM) exhibited strongly increased basal ATPase activities (3.3 – 5.4 ATP/min/monomer) that were further increased by substrate casein (1.64 – 2.47-fold stimulation) in contrast to ClpC WT (**Fig. 2A/B**). Casein-stimulated ClpC MD mutants reached 33% of total ATPase activity determined for MecA-activated ClpC WT. N-domain deletion also increased basal ATPase activity of ΔN-ClpC. Combining ΔN-ClpC with M-domain mutants had an additive stimulatory effect on ATPase activities, suggesting that the underlying mechanisms are distinct from one another (**Fig. 2A**). We infer increased basal ATPase activities of ClpC MD mutants can explain adaptor-independent substrate degradation. However, ATPase activation alone is not sufficient to explain ClpC activation as ΔN-ClpC did not allow degradation of GFP-SsrA despite having increased basal ATPase activity. This indicates specific consequences of MD mutations on ClpC conformation and activity.

**Figure 2.**
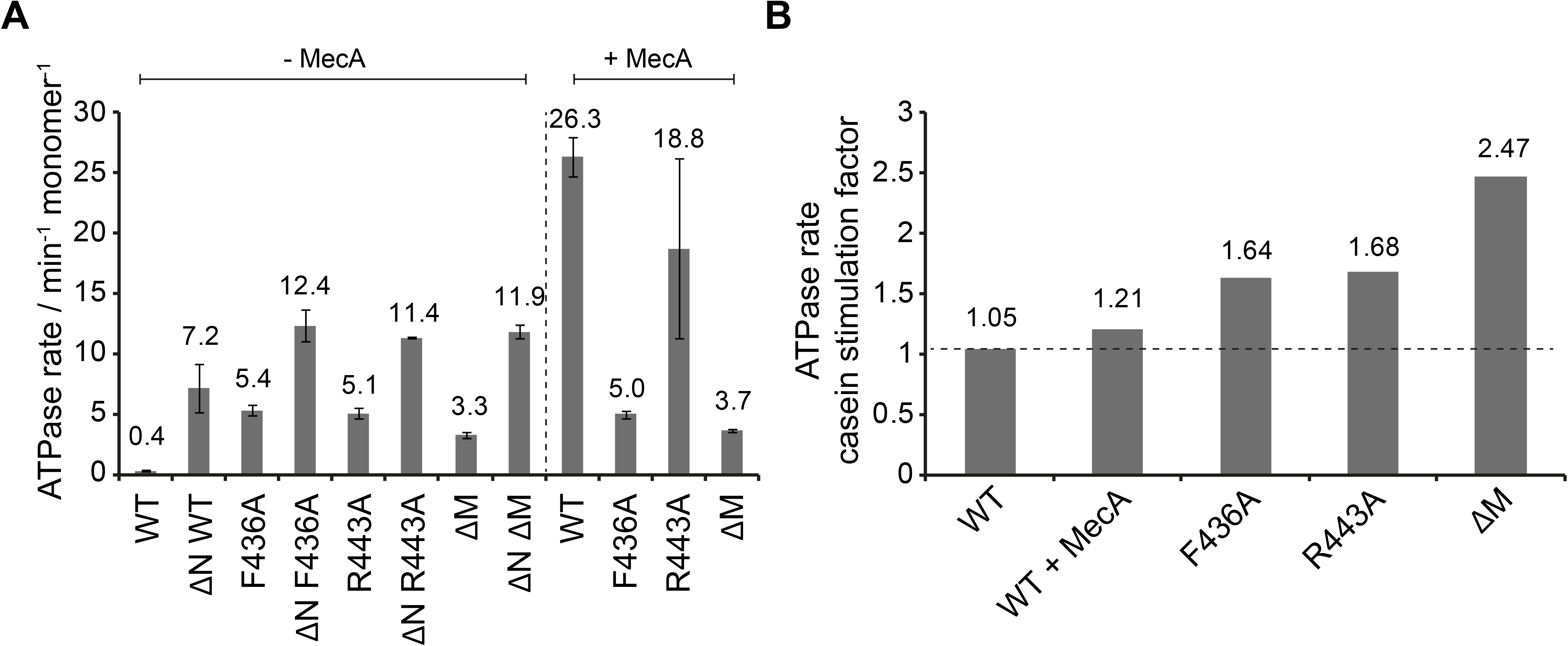
ClpC MD mutants exhibit increased basal ATPase activity that can be stimulated by substrate. (**A**) ATPase activities of ClpC wild type (WT) and indicated deletion variants (ΔN, ΔM) and MD mutants were determined in absence and presence of MecA. (**B**) ATPase activities of ClpC wild type (WT, ± MecA) and indicated MD mutants were determined in absence and presence of substrate casein. ATPase activities determined without casein were set as 1 and the relative increase of ATP hydrolysis in presence of casein was determined (stimulation factor).

### Head-to-head interactions of M-domains mediate formation of an inactive ClpC resting state

To understand the structural basis underlying the control of ClpC activity by the MD, we determined the cryo-EM structures of *S. aureus* ClpC with and without its activator MecA in the presence of ATPγS. The structure of ClpC on its own was solved at 8.4 Å resolution using ~ 90.000 collected particles (**Supplementary Fig. 4A**). Surprisingly, raw images and subsequent 2D classification revealed immediately that ClpC assumes a conformation different from the canonical hexameric arrangement of AAA+ proteins (**Supplementary Fig. 4B/C**). Attempts of 3D classification and refinement using the existing ClpC-MecA structures ^30,43^ failed to give a refined map, indicating that on its own, ClpC assumes a substantially different conformation. Indeed, reconstructions revealed that ClpC without MecA assembles into an almost decameric assembly made of two open spirals that interact via head-to-head contacts mainly mediated by the MDs (**Fig. 3A/B, Supplementary Fig. 4D**). Continuous spiraling of AAA+ proteins is often observed in X-ray structures ^17,18,44,45^ and ClpC half spirals are similar to these crystal packings. However, the ClpC double spiral is completely open on one side (**Fig. 3A/B, Supplementary Movie1**) giving a cradle-like molecule with the peripheral subunits more mobile than the core ones, as shown by lower local resolution (**Supplementary Fig 4E**). Additionally, high-threshold noise on the open part of the spiral indicates higher dynamics of this region, suggesting exchange of subunits. Accordingly, for peripheral ClpC subunits there is not sufficient density to account for all ClpC domains. The overall subdomain organization within the ClpC protomer is similar to that of the ClpC-MecA crystal structures, however domain positions and interactions are different. The AAA+ domains are staggered with a rise of ~ 20 Å per subunit (**Fig. 3B**). N-domains domains are packed between MDs and are displaced so that they lie on top of the adjacent small AAA1 subdomain (**Fig. 3B/C**). The MDs coiled-coils constitute the backbone of the spiral and mediate the spiral head-to-head contacts, which involve residues F436, R443 and D444 providing a structural rationale for the activated states of respective MD mutants (**Fig. 3D**).

**Figure 3.**
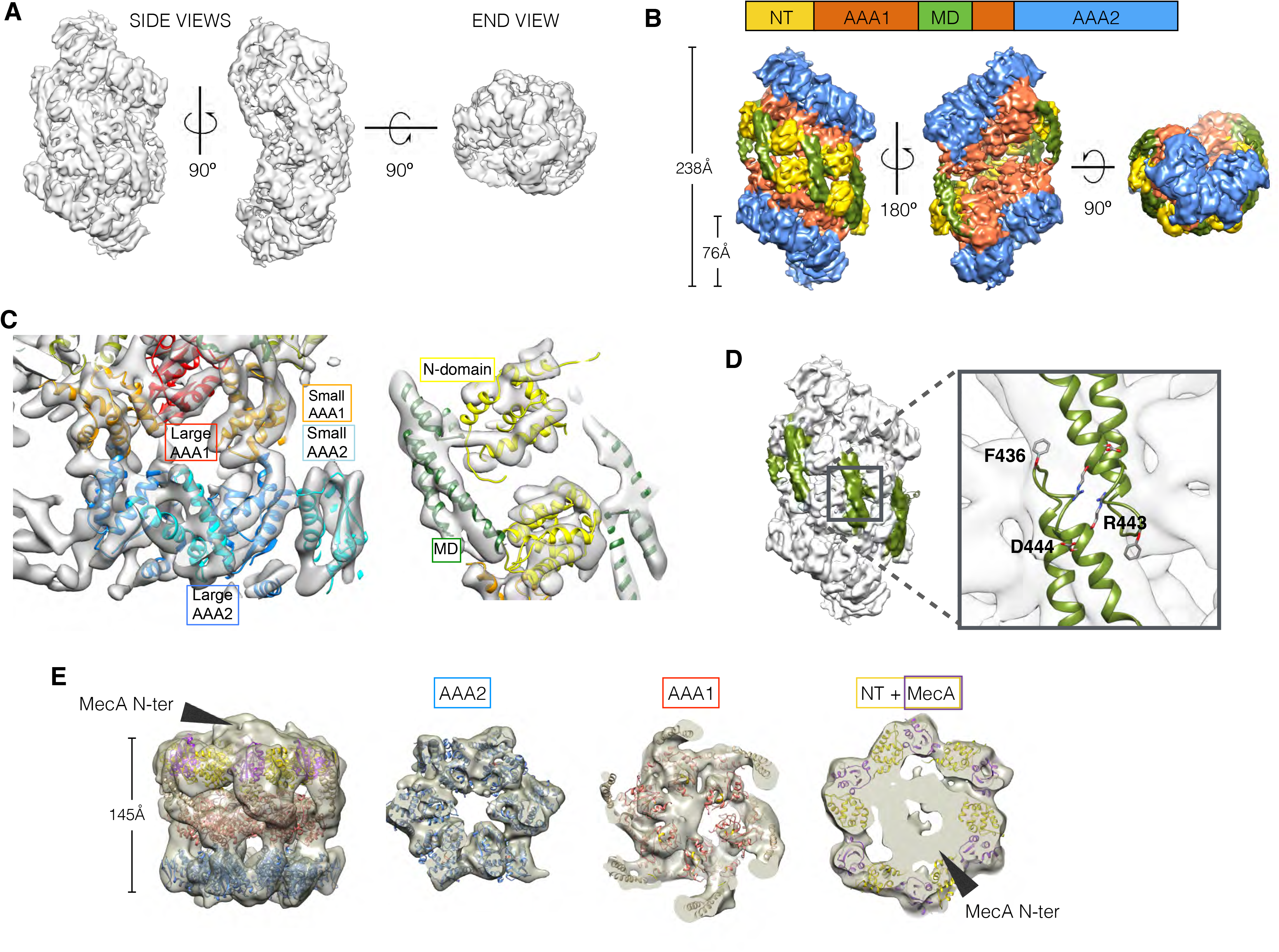
Cryo-EM structures of *S. aureus* ClpC-ATPγS with and without MecA. (**A**) Overview of ClpC 3D density map showing two open spirals that interact in a head-to-head manner. (**B**) ClpC density map coloured by domains. Head-to-head interactions between MDs (green) of each helical assembly are key contacts stabilizing the ClpC resting state. ClpC domain organization is given, including an N-terminal domain (NT), two AAA-domains (AAA-1, AAA-2) and the inserted coiled-coil M-domain (MD). (**C**) Details of fitting for each ClpC subdomain. (**D**) Zoomed view into MD-MD contacts highlighting conserved MD residues involved in interactions. (**E**) Structure of the ClpC-MecA complex. Fitted atomic model coloured by ClpC domains as in b and MecA C-terminal domains (purple). Density for MecA N-terminal domains is indicated.

We envision the helical ClpC assembly as a dynamic inactive resting state, which will interfere with substrate binding, association of the ClpP peptidase and interaction with adaptor proteins. Furthermore, the spiral arrangement of the AAA+ domains results in a displacement of most trans-acting arginine-fingers away from the adjacent nucleotide pocket (**Supplementary Fig. 4F**), thereby affecting ATP hydrolysis and explaining the low ATPase activity of ClpC WT.

The *S. aureus* ClpC-MecA cryo-EM map was reconstructed at 10 Å resolution with ~26,000 particles, (**Supplementary Fig. 5A-D**) and shows the classical hexameric assembly previously described ^30,43^, with MecA interacting with both the MD and the N-domain of ClpC. Additional extra density caps the hexamer and accounts for the N-terminal domain of MecA, whose structure is unknown. Automatic flexible fitting revealed that the hexamer is slightly asymmetric with 5 out of 6 subunits more defined (**Fig. 3E**), similar to the AAA+ protein Vps4 ^46^. From the resting to the MecA-bound state the N-domains undergo a 45° rotation that repositions the MecA-binding loop region from being blocked by the MD of the neighbouring subunit to be available and engaged in MecA binding (**Supplementary Fig. 5 E/F**). Additionally, binding of MecA to MDs breaks head-to-head MD contacts and is therefore expected to prevent formation of the ClpC resting state.

Taken together, our structures of ClpC alone or in complex with MecA show a dramatic reorganization from a helical resting state to a planar canonical AAA+ hexamer, explaining ClpC activation by the adaptor MecA. Intermolecular head-to-head contacts of MDs form the backbone of the ClpC resting state, qualifying the MD as crucial negative regulatory element consistent with MD mutant characterization.

### MecA abrogates head-to-head M-domain interactions

To demonstrate direct head-to-head contacts of ClpC MDs we introduced cysteine residues at the tip of the MD (E435C, E437C) to probe for site-specific disulfide crosslinking. These mutations were introduced into ClpC-C311T to avoid interference of the endogenous Cys311 residue. ClpC-C311T/E435C and ClpC-C311T/E437C were active in MecA-dependent protein degradation under reducing conditions (**Supplementary Fig. 6A**). Formation of crosslink products under oxidizing conditions that were fully reverted by addition of reducing agent was observed for ClpC-C311T/E437C but not for ClpC-C311T/E435C and the ClpC-C311T control (**Supplementary Fig. 6B**). This documents specificity of disulfide crosslinking and agrees well with the ClpC resting state model, showing E437 residues of two interacting M-domains are facing one another while E435 residues are oriented in opposite directions (**Supplementary Fig. 6B**). E437C disulfide crosslinking was most efficient in absence of nucleotide or presence of ADP and ATP, and less efficient in presence of ATPγS (**Supplementary Fig. 6C**). Importantly, ClpC-E437C crosslinking in presence of MecA was strongly reduced (**Fig. 4A**), consistent with MecA binding to MDs preventing head-to-head MD interactions. Similarly, crosslinking efficiency was reduced for ClpC-F436A/E437C, indicating a crucial contribution of F436 to intermolecular MD contacts (**Fig. 4B**), consistent with the ClpC WT cryo-EM structure.

**Figure 4.**
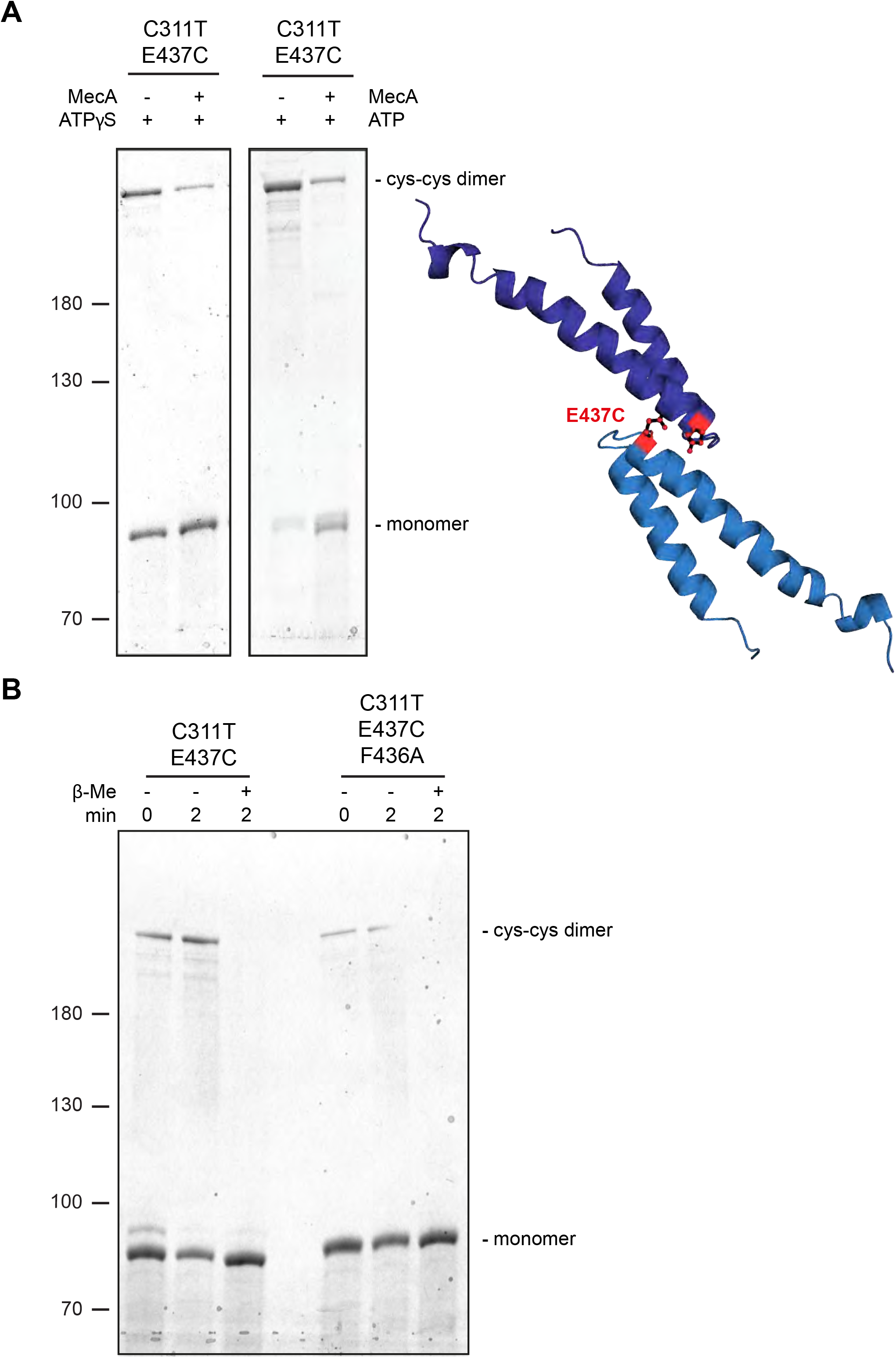
Disulfide crosslinking demonstrates MecA-sensitive head-to-head MD contacts. (**A**) Disulfide crosslinking of ClpC-C311T/E437C under oxidizing conditions was performed in presence of ATPγS or ATP without or with MecA and analyzed by subsequent non-reducing SDS-PAGE. A model of head-to-head interacting MDs is given and the position of E437 is indicated. (**b**) Disulfide crosslinking of ClpC- C311T/E437C and ClpC-C311T/F436A/E437C was performed in presence of ATPγS under oxidizing (+ Cu(Phe_3_)) and reducing (+β-mercaptoethanol) conditions and analyzed by subsequent non-reducing SDS-PAGE.

### Obstructing head-to-head M-domain contacts allows for ClpC hexamer formation

To provide biochemical support for the formation of a large, inactive ClpC resting state we employed chemical crosslinking and size exclusion chromatography. We used the ATPase-deficient ClpC-E280A/E618A variant (referred to as ClpC-DWB), harboring mutated Walker B motifs in both AAA+ domains allowing for ATP binding but not hydrolysis, facilitating analysis of adaptor or substrate impact on ClpC assembly. We first determined sizes of ClpC assemblies by glutaraldehyde crosslinking (**Fig. 5A**). ClpC-DWB was crosslinked to very large assemblies that were just entering the separating gel in SDS-PAGE in absence and presence of ATP (**Fig. 5A**). Addition of MecA allowed for formation of a smaller high molecular weight complex similar in size to crosslinked ClpB hexamers that were used as reference (**Fig. 5A**). Presence of MecA in the crosslinked ClpC-DWB complexes was confirmed by western-blot analysis (**Supplementary Fig. 7**). ClpC-F436A-DWB stayed monomeric in absence of nucleotide, indicating that large assemblies observed for ClpC-DWB rely entirely on MD contacts (**Fig. 5A**). ClpC-F436A-DWB crosslinking in presence of ATP caused formation of defined ClpC-F436A-DWB complexes that were similar in size to crosslinked ClpB hexamers. These findings confirm the predicted critical contribution of the MD to formation of a large resting state and the role of MecA in converting this assembly into a functional hexamer.

**Figure 5.**
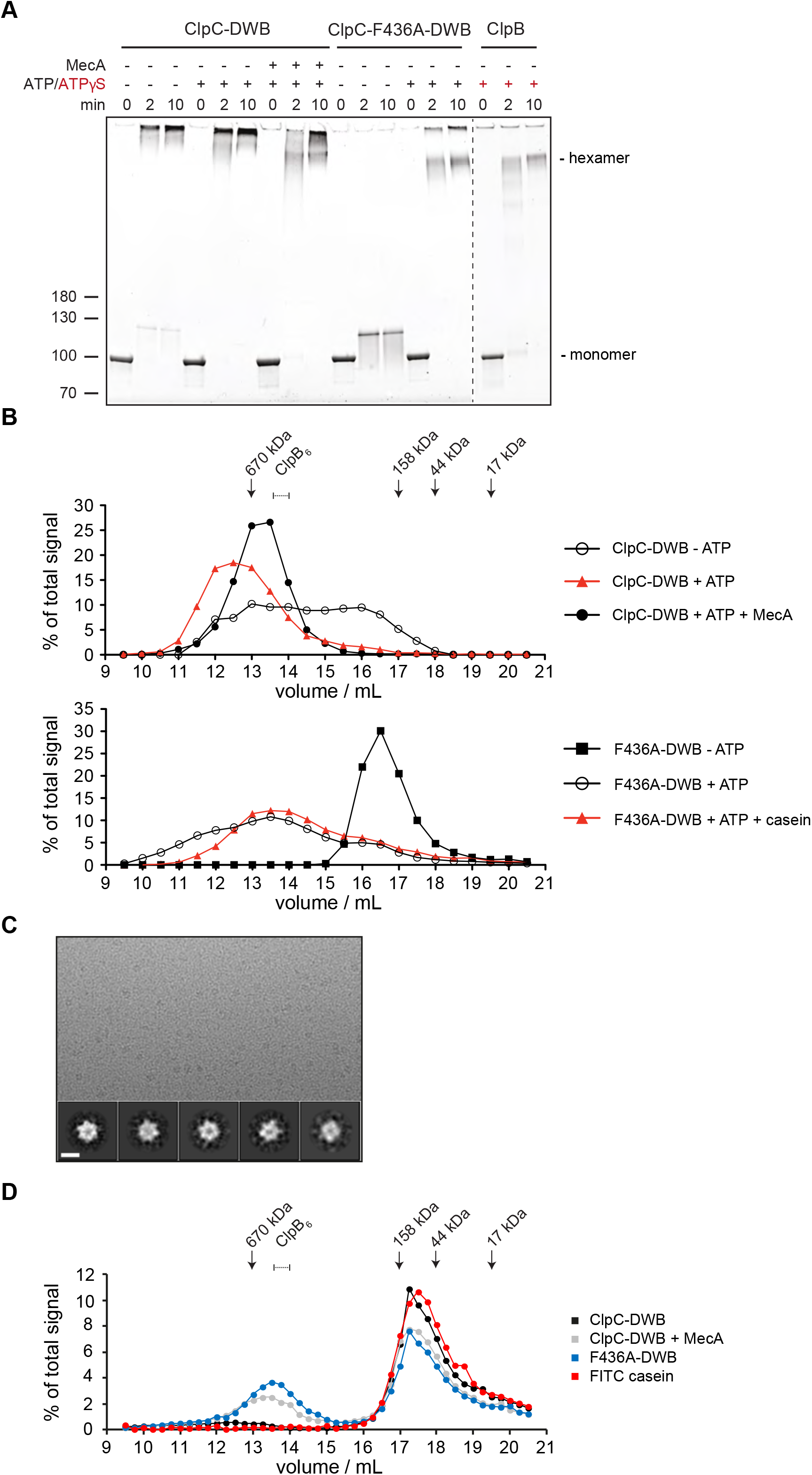
ClpC forms a large, inactive resting state that is sensitive to MecA and MD mutation. (**A**) Glutaraldehyde crosslinking of ClpC-E280A/E618A (ClpC-DWB) and a respective MD mutant variant (F436A) was performed in absence and presence of ATP without and with MecA as indicated. Crosslinking of *E. coli* ClpB in presence of ATPγS served as reference defining crosslinked hexameric assemblies. Crosslink products were analyzed by SDS-PAGE. (**B**) Oligomeric states of ClpC-DWB and ClpC-F436A-DWB were determined in absence and presence of ATP. Addition of MecA and casein is indicated. Elution fractions were analyzed by SDS-PAGE and quantified. Positions of peak fractions of a protein standard and ClpB-E279A/E678A hexamers (+ATP) are indicated. (**C**) Micrograph of ClpC-F436A sample in presence of casein (top). Examples of single particles are circled. 2D class averages of ClpC-F436A (bottom). Scale bar is 10 nm. (**D**) Binding of FITC-casein to ClpC-DWB and ClpC-F436A-DWB was analyzed in presence of ATP by size exclusion chromatography. FITC-casein fluorescence of elution fractions was quantified. Positions of peak fractions of a protein standard and ClpB-E279A/E678A hexamers (+ATP) are indicated.

We next analyzed sizes of ClpC complexes by size exclusion chromatography (**Fig. 5B, Supplementary Fig. 8A**). In absence of ATP ClpC-DWB showed a broad elution profile, suggesting formation of variable assemblies ranging from monomers to hexamers. ATP addition caused formation of larger ClpC assemblies that eluted prior to ClpB hexamers and a 670 kDa standard protein, suggesting formation of ClpC complexes larger than hexamers (**Fig. 5B, Supplementary Fig. 8A**). This was confirmed by static light scattering (SLS) measurements, revealing a molecular mass of ≈ 956 kDa corresponding to a decameric complex, consistent with cryoEM analysis (**Supplementary Fig. 8B**). Presence of MecA sharpened the ClpC elution profile and shifted ClpC-DWB fractions to later elution volumes right after the 670 kDa standard protein and now overlapping with ClpB hexamers (**Fig. 5B, Supplementary Fig. 8A**). Quantification of co-eluting MecA suggests the formation of a 1:1 ClpC:MecA complex, consistent with the binding stoichiometry determined for *B. subtilis* ClpC/MecA complexes and mass determination by SLS (767 kDa) (**Supplementary Fig. 8B**)^30,40^. This indicates that MecA binding shifts ClpC from a large, non-hexameric assembly to a ClpC6/MecA6 complex. We next analyzed the elution profile of ClpC-DWB-F436A. In absence of ATP the MD mutant eluted as defined species right at the elution volume of a 158 kDa standard protein suggesting formation of monomers/dimers (**Fig. 5B**). This indicates that the formation of larger assemblies noticed for ClpC WT (- ATP) depends on the M-domain, consistent with results from glutaraldehyde crosslinking (**Fig. 5A**) Unexpectedly, ClpC-DWB-F436A showed a broad elution profile upon ATP addition. While the ClpC-DWB-F436A peak fraction eluted right after the 670 kDa standard, larger assemblies at earlier elution volumes were also present. We speculated that activated ClpC MD mutants might be capable of recognizing themselves as substrate resulting in formation of larger complexes and explaining the elution profile. Indeed, we observed autodegradation of ClpC-F436A, ClpC-R443A and ClpC-ΔM but not ClpC WT in presence of ClpP (**Supplementary Fig. 9**). To prevent self-recognition of ClpC-DWB-F436A we repeated the size exclusion analysis in presence of substrate casein excess (**Fig. 5B**). Presence of casein sharpened the ClpC-DWB-F436A elution profile that was comparable to ClpC-DWB/MecA complexes and distinct from ClpC-DWB, suggesting hexamer formation, which was confirmed by mass determination by SLS (587 kDa). In contrast, casein addition did not cause formation of smaller ClpC-DWB complexes (**Supplementary Fig. 8B**). To further prove ClpC-F436A hexamer formation, a small cryo-EM dataset of ClpC-F436A with casein and ATPγS was collected and analyzed via 2D classification. Classes indicate that the ClpC-F436A assembles in an hexamer similar to ClpC WT with MecA (**Fig. 5C**).

Together our findings demonstrate that ClpC-DWB forms large, non-hexameric assemblies in a MD dependent manner, supporting the derived ClpC WT cryo-EM structure. This large ClpC-DWB assembly should neither allow for efficient substrate binding nor association with the ClpP peptidase. This prediction was confirmed by size exclusion chromatography showing poor interaction with ClpP and negligible binding to substrate FITC-casein (**Fig. 5D, Supplementary Fig. 8C**). In contrast, efficient binding to FITC-casein and ClpP was observed upon addition of MecA and for ClpC-DWB-F436A (- MecA) (**Fig. 5d, Supplementary Fig. 8C**). These findings were further confirmed by monitoring FITC-casein binding by anisotropy measurements (**Supplementary Fig. 8D/E**).

### M-domain activity control of ClpC is crucial for cellular viability

ClpC MD mutants allow for adaptor-independent and thus constitutive and uncontrolled ClpC activity. We wondered whether this loss of ClpC activity control has physiological consequences and co-expressed *S. aureus* ClpC WT, ΔN-ClpC and respective F436A mutants from an IPTG-regulated promoter together with *S. aureus* ClpP in *E. coli* cells. This strategy allowed us to only monitor potential toxic effects of ClpC M-domain mutants without interference by loss of endogenous functions of ClpC/MecA in *S. aureus* cells. Levels of ClpC variants were similar after 1 h of IPTG-induced protein production (**Supplementary Fig. 10A**). In case of ClpC-F436A we noticed accumulation of a degradation product upon ClpP coexpression, as also observed *in vitro* (**Supplementary Fig. 9**), suggesting autoprocessing. We observed strong toxicity upon expression of ΔN-ClpC-F436A at all temperatures tested while ClpC-F436A expression became lethal at 37°C and 40°C (**Fig. 6A**). Toxicity of ClpC MD mutants was much higher as compared to ClpC-WT and ΔN-ClpC, indicating an essential cellular need for ClpC repression by MDs. Furthermore, toxicity of ClpC MD mutants was dependent on coexpression of *S. aureus* ClpP, suggesting that uncontrolled protein degradation caused by constitutively activation results in cell death (**Supplementary Fig. 10B**).

**Figure 6.**
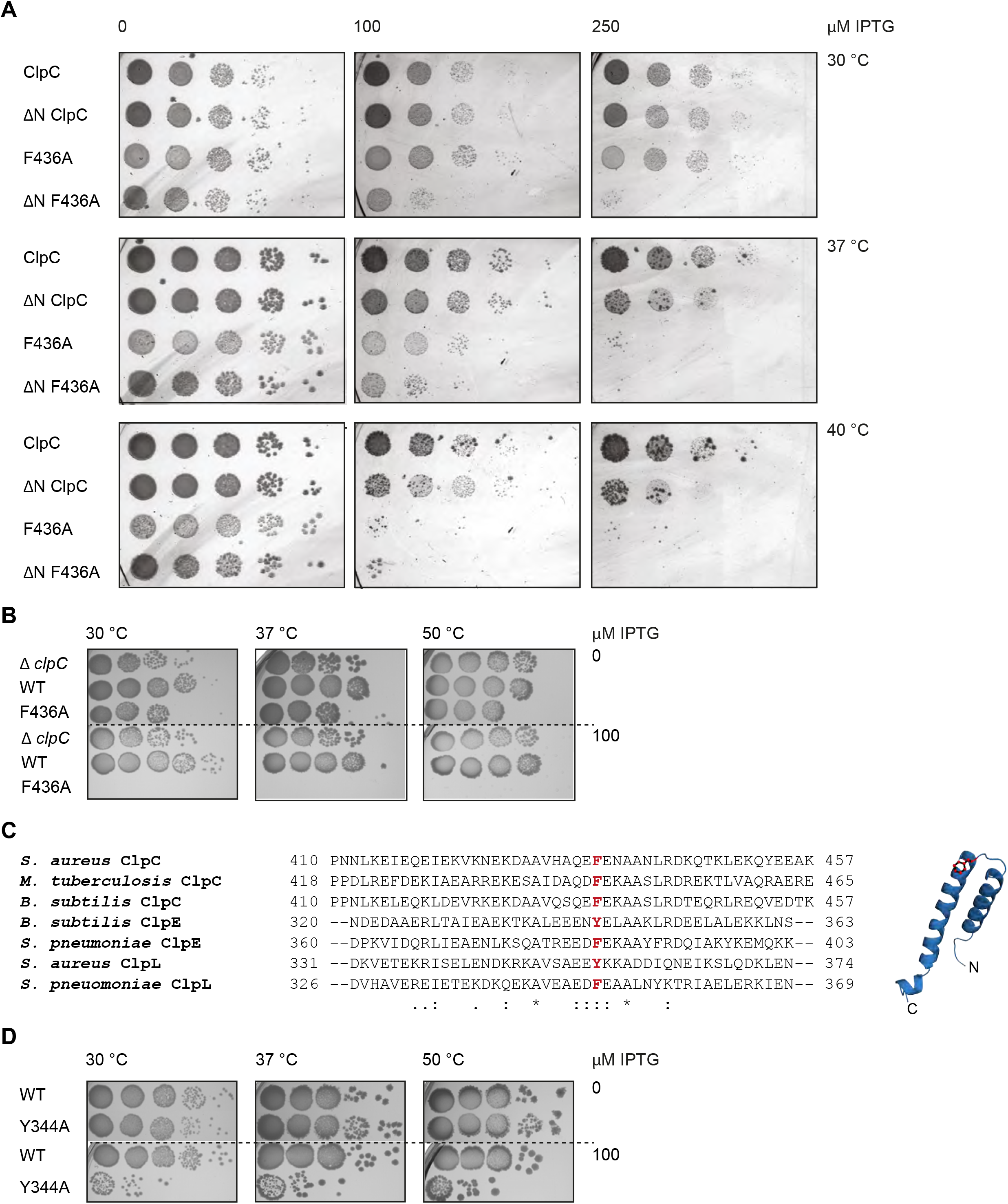
Loss of ClpC activity control is toxic *in vivo*. (**A**) *E. coli* cells constitutively expressing *S. aureus clpP* and harboring the indicated plasmid-encoded *clpC* alleles under control of an IPTG-regulatable promoter were grown overnight at 30°C and adjusted to OD600 of 1. Serial dilutions (10^-2^ – 10^-6^) were spotted on LB plates containing the indicated IPTG concentrations and incubated at 30°C, 37°C or 40°C for 24 h. (**B**) *B. subtilis* Δ*clpC* control cells and Δ*clpC* mutant cells harboring a *clpC* wild type (WT) or MD mutant (F436A) copy integrated at the *amyE*-locus under control of an IPTG-regulatable promoter were grown at 30°C to OD600 =1. Serial dilutions (10^-2^ – 10^-6^) were spotted on LB plates without or with 100 μM IPTG and incubated at 30°C, 37°C or 50°C for 24 h. (**C**) Sequence alignment of MDs from ClpC, ClpE and ClpL proteins. A highly conserved aromatic residue located at the tip of the coiled-coil structure is highlighted. (**D**) *B. subtilis* cells harboring an extra *clpE* wild type (WT) or MD mutant (Y344A) copy integrated at the *amyE*-locus under control of an IPTG-regulatable promoter were grown at 30°C to OD600 =1. Serial dilutions (10^-2^ – 10^-6^) were spotted on LB plates without or with 100 μM IPTG and incubated at 30°C, 37°C or 50°C for 24 h.

To further corroborate our findings we explored the physiological consequences of the same ClpC MD mutation in *B. subtilis*. Here, we deleted the chromosomal copy of the *clpC* gene (*clpC::tet*) and re-integrated either *clpC wt* or *clpC-F436A* at the *amyE*-locus under IPTG control. Expression of *clpC-F436A* but not *clpC wt* in presence of 100 μM IPTG was highly toxic at all temperatures (30°C – 50°C) (**Fig. 6B**). Levels of ClpC-wt and ClpC-F436A produced after IPTG addition in cells cultured in liquid medium were comparable, excluding differences in protein levels as reason for toxicity (**Supplementary Fig. 10C**). Importantly, growth of *clpC::tet* cells was not impaired, demonstrating that toxicity of ClpC-F436A reflects a gain-of-function phenotype and is not caused by loss of adaptor interaction (**Fig. 6B**). Together these findings demonstrate an essential role of MD mediated activity control for cellular viability.

The Hsp100 family members ClpE and ClpL harbor a coiled-coil MD that is similar in size to the ClpC MD and also displays some sequence homology. Notably, F436 and R443, identified here as key MD residues in ClpC activity control, are largely conserved in ClpE and ClpL M-domains (**Fig. 6C**). This suggests that the role of M-domains as crucial negative regulators of Hsp100 activity is conserved in other family members. To test for a conserved regulatory function we generated the *B. subtilis* ClpE-Y344A MD mutant corresponding to ClpC-F436A. *clpE wt* and *clpE-Y344A* copies were integrated at the *amyE*-locus in *B. subtilis* wild type cells under control of an IPTG-regulatable promoter. *B. subtilis* cells do hardly express endogenous *clpE* at non-stress conditions ^47,48^ thereby allowing mutant analysis without interference of ClpE WT copies. Expression of *clpE-Y344A* but not *clpE wt* in presence of 100 μM IPTG was highly toxic to *B. subtilis* cells at all temperatures tested (30°C – 50°C) (**Fig. 6D**). ClpE WT and ClpE-Y344A were produced to similar levels underscoring that toxicity is caused by deregulation of the ClpE MD mutant (**Supplementary Fig. 10D**). This suggests that ClpE MDs are also essential to downregulate ClpE activity preventing cellular toxicity.

## Discussion

In the presented work we established a new mechanism of activity control of ClpC, a central AAA+ chaperone widely distributed among Gram-positive bacteria. We show that coiled-coil MDs control ClpC activity in a unique manner by sequestering ClpC molecules in an inactive resting state.

Our findings extend the role of coiled-coil MDs as regulatory devices controlling AAA+ protein activity. MDs of the ClpB/Hsp104 disaggregases function as molecular toggles, which are crucial for AAA+ protein repression in the ground state and activation by an Hsp70 partner chaperone ^9,18-20,49^ (**Fig. 7**). Repression by ClpB/Hsp104 MDs relies on formation of a repressing belt around a canonical AAA+ ring by interacting with AAA-1 domains and neighboring MDs. Intermolecular head-to-tail contacts between long MDs (~ 120 residues forming two wings) are crucial to keep MDs in a horizontal, repressing conformation ^17^ (**Fig. 7**). Hsp70 binding to the tip of one wing breaks MD interactions and leads to ClpB activation.

**Figure 7.**
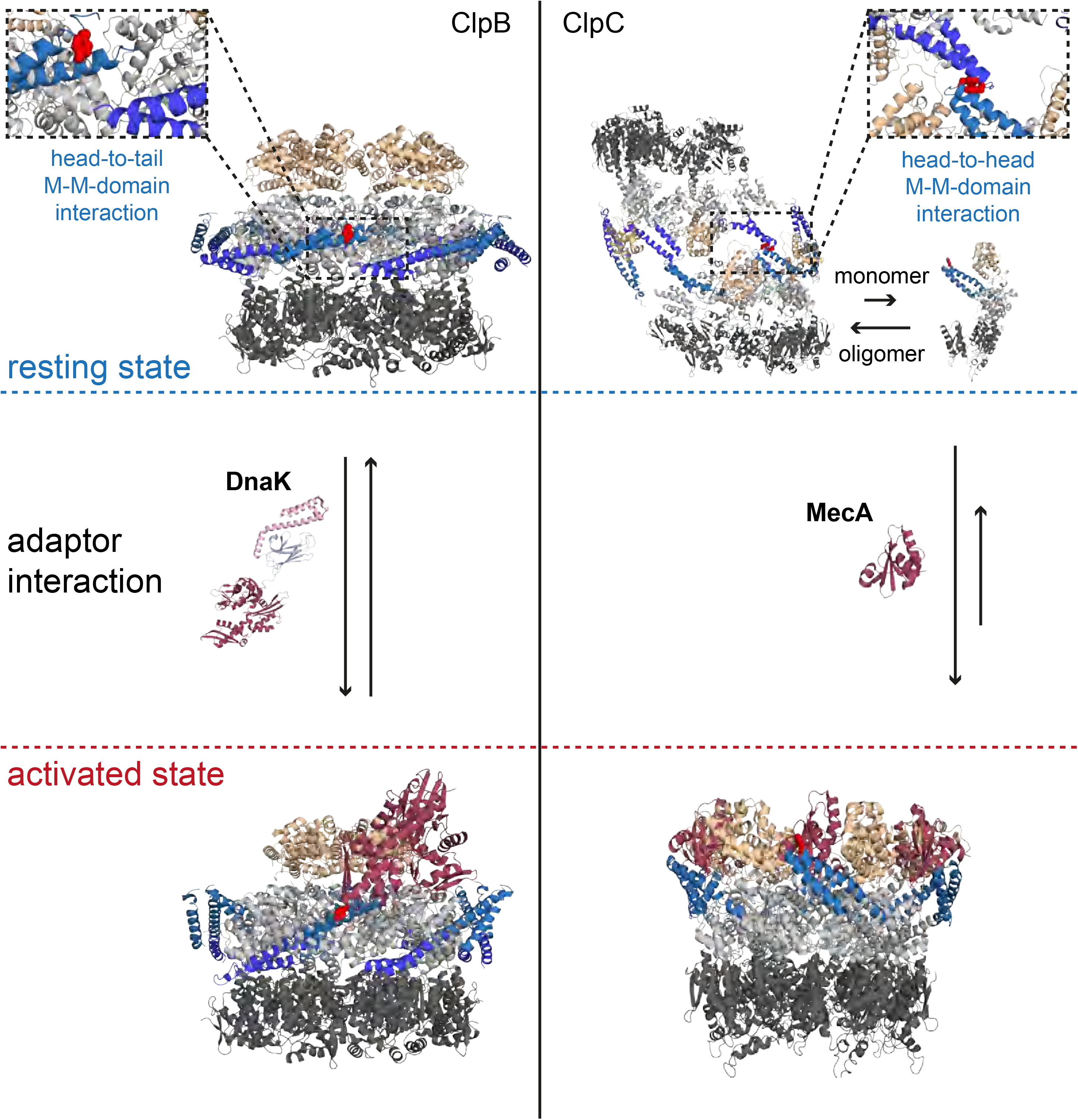
Regulatory coiled-coil MDs repress AAA+ protein activities by different mechanisms. ClpB is kept in a low activity resting state by long MDs forming a repressive belt around the hexameric AAA ring. MDs are kept in place by head-to-tail interactions between adjacent coiled-coils. In contrast, the ClpC resting state is formed by head-to-head MD contacts, allowing for assembly of two open ClpC spirals. Adaptor proteins of ClpB (DnaK) and ClpC (MecA) break MD contacts by binding to MD sites crucial for MD interactions. This results in AAA+ protein activation by releasing MD repression on ATPase activity (ClpB) or allowing for formation of active hexamers (ClpC).

ClpC MDs cannot function in the same manner due to their reduced size (~ 50 residues forming a single wing), which is too short to span the distance between neighboring subunits in a hexameric assembly. Instead, ClpC MDs form intermolecular head-to-head contacts allowing docking of two layers of ClpC molecules arranged in a helical conformation (**Fig. 7**). We define this large ClpC assembly as inactive resting state, as it strongly restricts binding of substrates, ClpP and adaptor proteins and does not allow for efficient ATP hydrolysis. The interaction surface of MDs is limited (50 Å^2^) suggesting that the ClpC storage state is not stable but dynamic, in agreement with our structural and biochemical analysis. This will open the route for ClpC activation upon dissociation of ClpC molecules from the resting structure, liberating MDs and N-domains for binding to activating MecA or other adaptors. Additionally, MecA might bind to peripheral subunits of the ClpC storage state causing their displacement. ClpC MDs thereby function as molecular switches, similar to ClpB/Hsp104 MDs, ensuring repression in the ground state and allowing for activation in presence of substrate recruiting adaptors (**Fig. 7**). This dual activity is best illustrated for MD residue F436 located at the tip of the coiled-coil structure. F436 is essential for both, intermolecular MD interaction and MecA binding (**Fig. 7**). The N-terminal domain appears to play an additional role in this switch of conformations, by going from a more hidden position in between MDs in the resting state to a more exposed one, available to MecA or other adaptors.

Our newly derived model of *S. aureus* ClpC activity control differs from a former one, showing that *B. subtilis* ClpC is monomeric and requires MecA for hexamer formation ^40^. We realize that former ClpC analysis was performed in presence of high salt concentrations (300 mM NaCl), which likely interfere with MD head-to-head contacts involving charged residues (R443, D444). A regulatory model involving only monomer-hexamer transitions cannot explain activation and severe cellular toxicity of ClpC MD mutants shown here in *E. coli* and *B. subtilis* cells. However, we suggest that the oligomerization dependence of ClpC on MecA at high salt concentrations might represent a fail-safe system ensuring adaptor-dependent ClpC activation under conditions that do not allow for resting state formation (e.g. salt stress).

Once formed the hexameric ClpC_6_/MecA_6_ complex is stable and does not dissociate spontaneously, raising the question how ClpC activation is turned off. Adaptor proteins are targeting themselves for autodegradation by ClpC/ClpP if substrates are no longer available ^15,29,50^. This mechanism couples substrate availability with ClpC activation and ensures fast ClpC inactivation in absence of substrate by causing dissociation of ClpC hexamers into monomers that are subsequently sequestered in the resting state. Notably, other chaperone machineries including the Hsp70 member BIP ^51^ and the AAA+ protein Rca ^52^ also form large, inactive resting states that are converted into active species depending on substrate availability. Sequestration of chaperones therefore seems a more widespread activity to tune their activities according to the physiological need.

Constitutively activated ClpC MD mutants exert strong toxicity in *E. coli* and *B. subtilis* cells, demonstrating an essential physiological need to tightly control ClpC function. We assume that overhasty protein degradation by deregulated ClpC/ClpP complexes of e.g. newly synthesized proteins or secretory proteins, which did not yet reach their native structures or cellular compartment, leads to cell death.

Homologous MDs are present in the AAA+ protein family members ClpE and ClpL and key residues driving formation of the ClpC resting state are evolutionary conserved. We show that mutating a key regulatory MD residue of ClpE also causes cellular toxicity, strongly suggesting that the repressing mode of MDs established here for ClpC is also operational in ClpE and ClpL and thus is a more general mechanism for controlling bacterial AAA+ chaperone systems.

Severe toxicity of ClpC and ClpE MD mutants qualifies these AAA+ chaperones as targets for antimicrobials. In fact, the deregulation of bacterial proteases represents a novel antibacterial strategy ^53,54^. Notably, the *M. tuberculosis* ClpC N-terminal domain was recently identified as target of cyclic peptides with antibacterial activities ^55-57^. Although the mode of these drugs and their effects on ClpC function are not understood it is likely that they interfere with ClpC activity control. Our findings presented here open a new route for toxic ClpC deregulation by identifying MDs as crucial regulatory elements and thus drug targets and offering a promising approach to attack multi-drug resistant bacteria.

## Materials and Methods

### Strains, plasmids and proteins

*E. coli* strains used were derivatives of MC4100, XL1-blue or DH5α. ClpC, MecA, ClpP were amplified by PCR, inserted into pDS56 and verified by sequencing. Mutant derivatives of ClpC were generated by PCR mutagenesis and standard cloning techniques in pDS56 and were verified by sequencing. Transformation into *B. subtilis* 168 was performed by standard methods ^58^. AmyE insertion in *B. subtilis* was checked by plating on agar containing 0.4 % starch (w/v) additionally to appropriate antibiotics, screening for successful loss of α-amylase by staining starch with Lugol’s iodine.

ClpC and variants, MecA and ClpP were purified after overproduction from *E. coli* Δ*clpB::kan* cells. GFP-SsrA was purified after overproduction from *E. coli* Δ*clpX* Δ*clpP* cells. All proteins were purified using Ni-IDA (Macherey-Nagel) and size exclusion chromatography (Superdex S200, GE Healthcare) following standard protocols. Pyruvate kinase of rabbit muscle, casein and FITC-casein were purchased from Sigma. Protein concentrations were determined with the Bio-Rad Bradford assay.

### Biochemical assays

#### Size exclusion chromatography and multi-angle light scattering

Complex formation of ClpC (10 μM) was monitored by size exclusion chromatography (SEC, Superose 6 10/300 GL, GE Healthcare). MecA (20 μM), ClpP (20 μM), casein (16.66 μM) and FITC-casein (2.5 μM) were added as indicated. Experiments were run at 25°C in buffer A (50 mM Tris pH 7.5, 25 mM KCl, 20 mM MgCl_2_) supplemented with 2 mM ATP. Samples were prepared freshly and incubated for 5 minutes with 2 mM ATP prior to injection. Fractions were collected in 96-well plates, aliquots taken and subjected to SDS-PAGE. Gels were stained using SYPRO® Ruby Protein Gel Stain (ThermoFisher) following manufacturer instructions. Band intensities were quantified using ImageJ. To monitor the binding of ClpC to FITC-Casein, the collected fractions were analyzed for FITC fluorescence using FLUOstar Omega (BMG Labtech) with standard FITC filter sets. Chromatography was performed in three independent experiments each and representative results are provided.

Complex formation analyzed by SEC was additionally followed by online multi-angle light scattering (MALS) using an Agilent 1260 Infinity II HPLC system connected in series with a 3-angle multi-angle light scattering detector (miniDAWN TREOS II, Wyatt Technology, collection rate of 2 data points per second) and an additional online differential refractive index detector (Optilab T-rEX, Wyatt Technology) for concentration determination. Data analysis was performed using ASTRA 7.1 (Wyatt Technology). Samples were additionally filtered through a 0.2 µm low-protein binding syringe filter (Millex-GV, Merck Millipore Ltd.) before application to the SEC-column.

#### ATPase activity

The ATPase rate of ClpC and mutants was determined using a coupled-colorimetric assay as described before ^9^. The assay was carried out at 2 mM ATP in buffer A including 2 mM DTT at 30°C using a FLUOstar Omega plate reader. The final protein concentrations were as follows: ClpC (1 µM), MecA (1.5 µM), casein (10 µM). In presence of casein MecA concentrations were reduced to 0.2 µM. The raw data was analyzed using the following equation:

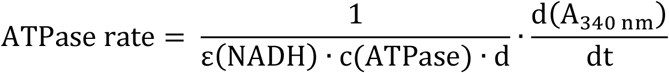

**ε (NADH):** Extinction coefficient at 340 nm for NADH (M^-1^ cm^-1^)

**c(ATPase):** Concentration of ATPase (M)

**d:** path length (cm)

**d(A340 nm)/dt:** derivative of the linear graph (slope)

ATPase rates were calculated from the linear decrease of A340 in at least three independent experiments and standard deviations were calculated.

#### Degradation assays

FITC-casein degradation was analyzed using a CLARIOstar plate reader, in black 384 well plates (Corning, NBS coated, flat bottom), in buffer A with 2 mM DTT. The final protein concentrations were as follows 0.3 µM FITC-casein, 1 µM ClpC, 1.5 µM MecA, 2 µM ClpP. The assay was carried out in the presence of an ATP regenerating system (0.02 mg/mL PK, 3 mM PEP pH 7.5) and 2 mM ATP. For measuring the decrease of FITC-casein fluorescence the filters 483-14 nm (ex) and 530-30 nm (em) were used. For data processing the background in the absence of ClpC was subtracted and the initial fluorescence intensities were set to 1. FITC-casein degradation rates were determined by the initial slopes of the fluorescence signal increase in at least three independent experiments and standard deviations were calculated. Alternatively degradation of FITC-casein (5 µM) was monitored by SDS-PAGE followed by Coomassie staining.

Degradation of GFP-SsrA (0.2 µM) was performed in buffer A with 2 mM DTT using the following protein concentrations: 1 µM ClpC, 1.5 µM MecA, 2 µM ClpP. Reactions were started by addition of an ATP regenerating system (0.02 mg/mL PK, 3 mM PEP pH 7.5) and 2 mM ATP. GFP fluorescence was monitored with a LS55 spectrofluorimeter (Perkin Elmer) using 400 and 510 nm as excitation and emission wavelengths. Initial GFP-SsrA fluorescence intensity was set as 100 and degradation rates were determined by the initial slopes of fluorescence signal decrease in at least three independent experiments and standard deviations were calculated.

#### Crosslinking

Glutaraldehyde crosslinking was performed by incubating 1 µM ClpC or ClpB buffer B (50 mM HEPES, 25 mM KCL, 10 mM MgCl_2_, 2 mM DTT, pH 7.5) in absence or presence of 2 mM ATP/ATPγS and 3 µM MecA at 25°C for 15 minutes. Crosslinking was started by adding Glutaraldehyde (Sigma) to a final concentration of 0.1%. Aliquots were taken at indicated time points and crosslinking was quenched by adding Tris (pH 7.5) to a final concentration of 50 mM. Samples were subjected to SDS-PAGE and gels stained with SYPRO® Ruby Protein Gel Stain (ThermoFisher).

Disulfide crosslinking was performed by incubating 3 µM ClpC in buffer A (in the absence or presence of 2 mM nucleotide). MecA (4.5 µM) was added as indicated MecA and 2 mM β-Mercaptoethanol at 25°C for 5 minutes. Crosslinking was started by addition of copper-phenanthroline to a final concentration of 100 µM. Aliquots were taken and crosslinking was stopped by adding SDS sample buffer without β-mercaptoethanol but containing 4 mM iodacetamide. Samples were boiled and analyzed by SDS-PAGE followed by Coomassie staining.

Crosslinking was performed in three independent experiments each and representative results are provided.

#### Anisotropy measurements

FITC-casein (100 nM) was incubated with varying concentrations ClpC in buffer A with 2 mM DTT for 1 h in absence or presence of 2 mM ATPγS. Changes in fluorescence polarization were determined using a CLARIOstar plate reader (BMG Labtech) at 482 and 530 nm excitation and emission wavelengths (Target mP 35).

#### Western blotting

SDS-PAGEs were transferred to nitrocellulose or PVDF membranes by semi-dry blotting or wet blot transfer. Membranes were subsequently blocked with either 3% BSA (w/v) or 5% (w/v) skim milk powder in TBS-T. Custom-made antibodies were used at the following dilutions: anti-ClpC (*B. subtilis*) 1:100.000, anti-ClpE 1:30.000, anti-ClpC (S. aureus) 1:50.000 and anti-MecA 1:30.000. anti-rabbit alkaline phosphatase conjugate (Vector Laboratories) was used as secondary antibody (1:10.000). Blots were developed using NBT/BCIP or ECF™ Substrate (GE Healthcare) as reagent and imaged via Image-Reader LAS-4000 (Fujifilm). Western blotting was performed in three independent experiments each and representative results are provided.

### ClpC structure determination by cryo-electron microscopy

#### Cryo-electron microscopy

*S. aureus* ClpC WT (6 µM) was incubated for 15 min at room temperature in 25 mM Tris-HCl (pH 7.5), 25 mM KCl, 10 mM MgCl_2_, 1 mM DTT and 2 mM ATPγS. For ClpC-MecA complex formation, *S. aureus* MecA was incubated with ClpC in a 3:1 molar ratio. Samples were vitrified with liquid ethane on Quantifoil R2/2 grids using a Vitrobot Mark IV (FEI) at 100% humidity, 24°C temperature and blotting time of 3 seconds.

Images of ClpC were collected using the EPU software on a Titan Krios TEM (FEI) operating at 300kV, using a Falcon 2 direct electron detector (FEI). Images of ClpC in complex with MecA were collected using the EPU software on a Titan Krios TEM (FEI) operating at 300kV equipped with a Gatan K2 Summit direct electron detector and bioquantum energy filter with 20 eV slit. The defocus range was set between -1 and -3 µm with a total dose of 30 electrons/Å^2^ in 17 frames for ClpC and 50 electrons/Å^2^ in 40 frames for ClpC-MecA. Pixel size was 1.34 Å/pixel for ClpC and 1.37 Å/pixel for ClpC-MecA.

#### Image processing

Movie frames alignment with dose weighting ^59^ and CTF estimation ^60^ was performed on-the-fly using a Scipion suite ^61^. ClpC particles were picked with Gautomatch and a dataset of ~90.000 particles from 1100 micrographs was generated. ClpC-MecA complex particles were picked using Gaussian picking in RELION ^62^ and ~500.000 particles from 2100 micrographs were obtained. The initial datasets were subjected to reference-free 2D classification in order to clean the datasets.

Initially, for 3D processing of ClpC (**Supplementary Fig**. **4**), the crystal structure of ClpC-MecA ^30^ low-pass filtered at 60 Å was used, but the dataset failed to refine. Attempts to use *ab initio* models generated with Eman2 ^63^ also did not result in high-resolution 3D refinement. As the ClpC assembly appeared much larger in size than the ClpC-MecA hexamer, we generated a cylindrical starting model by filtering to 70 Å two copies of ClpC-MecA stuck back to back with the AAA2 rings in contact. With this starting model ClpC refined to 8.4 Å resolution as estimated with the 0.143 FSC criterion, with visible separated helices. The same result was confirmed by generating an *ab-initio* starting model using the SGD method implemented in cryoSPARC ^64^ and refining the structure within the same program suite. Reconstructions with and without C2 symmetry applied were performed (**Supplementary Fig**. **4, g-h)** to a similar resolution and the C2 map was used for display as it allows a better visualization of N-domains.

For 3D processing of ClpC-MecA (**Supplementary Fig. 5**) the crystal structure of the *B. subtilis* ClpC-MecA complex filtered at 60Å was used (pdb code: 3PXI). A large dataset was initially used, but particles were preferentially oriented so a reduced dataset (~30.000 particles) with balanced angular distribution was used to reduce anisotropy. Both asymmetric and six fold symmetric maps were built. Even though the nominal resolution of the symmetrized map was better, the reconstruction appeared over filtered, thus indicating that artifacts were caused by forced symmetrisation. The final map was reconstructed at 11 Å resolution. Local resolution was evaluated using the local resolution tool of RELION.

#### Model building and fitting

Models of *S. aureus* ClpC and MecA were generated using Phyre ^65^. The N-domain to AAA1 link was refined using the Chimera loops modeller tool ^66^. Initial manual fitting was performed and then automatically refines using Imodfit ^67^.

### Spot tests

*E. coli* cells harboring plasmid-encoded *clpC* alleles were grown in the absence of IPTG overnight at 30°C. Serial dilutions were prepared, spotted on LB-plates containing different IPTG concentrations and incubated for 24 h at indicated temperatures. *B. subtilis* strains were inoculated with a fresh overnight culture to an OD_<u>600</u>_ of 0.05 and grown to mid-exponential growth phase. Optical densities of all strains were adjusted to OD_600_ of 1, serial dilutions were performed and 10 µl (10^-2^ - 10^-6^) were dropped on agar plates (without or with 100 µM IPTG) and incubated overnight at indicated temperatures. Spot tests were performed in three independent experiments each and representative results are provided.

## Acknowledgments

K. B. F. was supported by the Hartmut Hoffmann-Berling International Graduate School of Molecular and Cellular Biology (HBIGS). This work was funded by grants of the Deutsche Forschungsgemeinschaft (BB617/17-2 and MO 970/4-2) to B.B. and A.M. and a fellowship of the Hannover School for Biomolecular Drug Research to I.H. and the DFG grants Tu106/8-1 and Tu106/6-2 to K.T. The Cryo-EM facility at the Science for Life Laboratory Stockholm University (M.C.) is supported by grants from the Knut and Alice Wallenberg Foundation and the Family Erling Persson Foundation. We thank Stefan Fleischmann for IT support, Christos Savva for microscopy support, Björn Forsberg and Shintaro Aibara for image-processing discussions, Helen Saibil for support in the early stage of the project and Armgard Janczikowski for technical assistance.

## Author contributions

Conceived and designed experiments: M.C., K.B.F., M.M., J.J., I.H., F.G., D.L., S.G., K.T., B.B., A.M. Performed experiments: M.C., K.B.F., M.M., J.J., I.H., F.G., D.L., S.G. Analyzed the data: M.C., K.B.F., M.M., J.J., I.H., F.G., D.L., S.G., K.T., B.B., A.M. Wrote the manuscript: M.C., B.B., A.M.

